# Prevalence of Group II Introns in Phage Genomes

**DOI:** 10.1101/2025.05.22.655115

**Authors:** Liana N. Merk, Thomas A. Jones, Sean R. Eddy

**Affiliations:** Department of Molecular and Cellular Biology, Harvard University, 02138, Cambridge, USA; Howard Hughes Medical Institute, Harvard University, 02138, Cambridge, USA; Harvard Graduate Program in Biophysics, Harvard University, 02138, Cambridge, USA

## Abstract

Although bacteriophage genomes are under strong selective pressure for high coding density, they are still frequently invaded by mobile genetic elements (MGEs). Group II introns are MGEs that reduce host burden by autocatalytically splicing out of RNA before translation. While widely known in bacterial, archaeal, and eukaryotic organellar genomes, group II introns have been considered absent in phage. Identifying group II introns in genome sequences has previously been challenging because of their lack of primary sequence similarity. Advances in RNA structure-based homology searches using covariance models has provided the ability to identify the conserved secondary structures of group II introns. Here, we discover that group II introns are widely prevalent in phages from diverse phylogenetic backgrounds, from endosymbiont phage to jumbophage.

## Introduction

Group II introns are self-splicing ribozymes capable of retromobility (1). First identified in fungal mitochondrial genomes (2; 3), group II introns have since been identified in all three domains of life: bacteria, archaea, and eukaryotic organelles (4). They are notably absent in eukaryotic nuclear genomes, though they are likely the ancestral progenitors of spliceosomal introns (1). Furthermore, despite their wide dispersal in bacterial genomes (5; 6), group II introns have been considered to be absent in phage (7). Only two examples have been mentioned in passing (8; 9). Group I introns, in contrast, have long been known in phage genomes and have been studied in depth (10). One explanation for the apparent disparity between the prevalence of group I introns and the lack of group II introns in phage could be simply that group II introns do exist in phage genomes, but have not been found yet.

Group II introns are challenging to identify by computational sequence analysis. They have little primary sequence conservation but strong secondary structure conservation. The consensus group II secondary structure consists of six domains called D1-D6 (Figure 1A). There is often an open reading frame (ORF) for an intron-encoded protein (IEP), typically in the D4 loop. ORF-less group II introns are typically around 600 nucleotides in length, and ORF-containing introns are typically around 2-3kb (11). The start of the D5 stem contains the catalytic triad (usually AGC or CGC) which coordinates metal ions in the splicing mechanism (4). The D6 loop contains a bulged adenosine that is the 2’-5’ branch point in the splicing mechanism. Because D5 and D6 contain the required sequences and structures required for the catalytic splicing mechanism, D5 and D6 are the most conserved part of the group II consensus structure. Previous computational efforts to identify new group II introns have focused on these conserved domains (12).

**Fig. 1:**
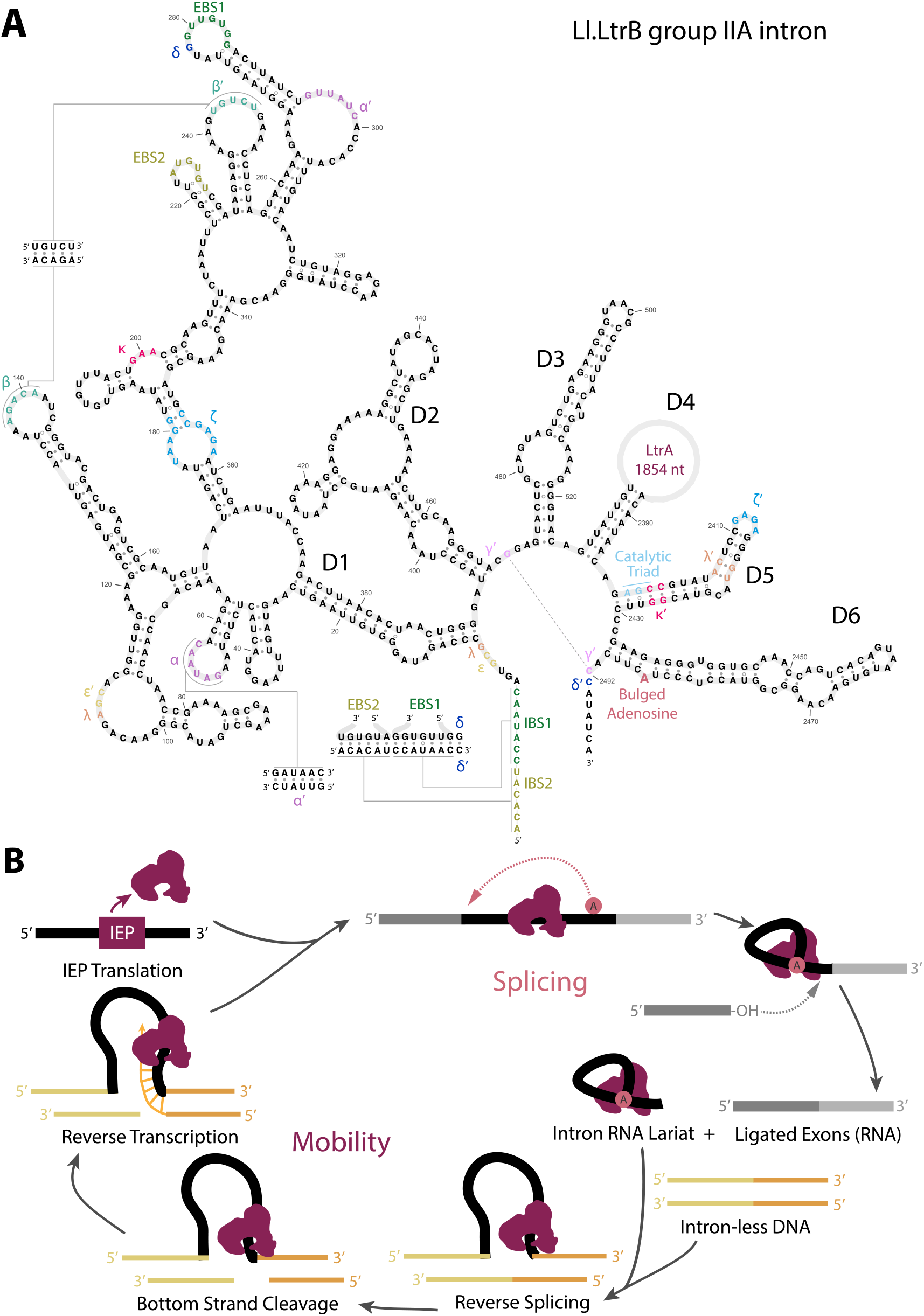
Group II intron consensus secondary structure and mechanisms. **A**. Secondary structure of *Lactococcus lactis* Ll.LtrB group II intron reproduced from D’Souza *et al*. (50). The LtrA IEP ORF is located within D4, shown as an open circle. **B**. Splicing and mobility mechanisms, adapted from (51). RNA is shown in grey and black, and DNA is shown in yellow and orange. The intron encoded protein (IEP) binds to intron RNA, enabling its folding into splicing-enabled structure. Once the lariat invades intronless DNA, the IEP begins target-primed reverse transcription.

Group II introns insert themselves into the DNA of an intronless host gene allele using a “retro-homing” mechanism catalyzed by the IEP. A group II IEP typically contains a reverse transcriptase (RVT) domain, a maturase domain, and an endonuclease domain (Figure 1B). Homing specificity is determined by base-pairing between intron sequences in D1 and exon sequences at the insertion site, called intron binding site (IBS) and exon binding site (EBS) sequences. Group II introns also transpose to new sites using the same mechanism, when a new site has sufficient IBS-EBS complementarity. Once the intron lariat RNA invades the homing site, the bottom DNA strand is nicked around 10 nucleotides downstream of the insertion site by the endonuclease, forming the primer needed for reverse transcription. Some group II introns only home during replication, using the transient ssDNA as a primer, in which case the IEP does not include an endonuclease domain (13). Previous computational search tactics have taken advantage of the fact that IEPs co-evolve with their group II introns (14), and IEPs can be identified by sequence homology. However, relying on IEP homology misses group II introns that are ORF-less or contain non-canonical IEPs, and conversely, there are many reverse transcriptase homologs that are not associated with group II introns.

A general computational method for identifying homologs of a conserved RNA secondary structure and sequence consensus uses probability models called profile stochastic context-free grammars (profile SCFGs, also called covariance models) (15; 16). A software package called Infernal implements profile SCFG-based search and alignment (17). The Rfam database of 4000+ known conserved RNA structure elements is built with Infernal (18). The input to an Infernal search is a multiple sequence alignment of the conserved RNA annotated with its consensus secondary structure. From this sequence and structure information, Infernal builds a consensus statistical model, which can then be used to search genome sequences for homologs, and to structurally align new homologs to the consensus. Profile SCFG algorithms used to be prohibitively computationally expensive, but a set of accelerated algorithms in Infernal now allows for comprehensive searches for RNAs in large genome and metagenome datasets, including RNAs the size of catalytic introns (19). Here we use Infernal to search for group II introns in phage genomes.

## Materials and Methods

Profile SCFGs for group II introns (RF00029 and CL00102) were from Rfam 14.10 database (18). The Millard dataset of 29,015 curated phage genomes was obtained with the INPHARED Perl script (20) on 14 Dec 2023, and phylogenetic metadata was updated on 11 Sept 2024. The IMG/VR metagenomic dataset was version 4.1 (21). Phage genome annotation to aid in determining insertion sites was performed with Bakta 1.9.1 (22) and Pharokka 1.5.1 (23).

Infernal searches used cmsearch from Infernal v1.1.4 (Dec 2020) with an E-value threshold of 0.01. Unannotated intron-encoded proteins were identified by translating within the intron bounds in three frames, then using hmmscan from HMMER 3.3.2 (24) against Pfam-A 35.0 (25) with an E-value cutoff of 10^−3^.

Multiple sequence alignments and phylogenetic tree inference for the reverse transcriptase domain and terminase large subunit (TerL) was done as follows. Profile hidden Markov models (profile HMMs) for RVT 1 (PF00078) and terminase large subunit (PF03237) were from Pfam 37.0. For RVT, PF00078 was used as an hmmsearch query to identify and align RVT domains from our seven intron IEPs to a set of annotated RVT domain sequences from a dataset from Toro *et al*. (26), randomly subsampling a maximum of 20 RVT domain sequences from each of the five classes (GII, DGR, G2L, Retron-like, RT/CRISPR-Cas) defined by Toro *et al*.. For TerL, PF03237 was used to identify TerL homologs in an iterative two-round hmmsearch of all Millard phage genomes, plus an additional single hmmsearch of a subset of intron-containing IMG/VR genomes that contained a D1-D4 hit and D5/D6 hit within 3 kb of each other. TerL hits in the Millard dataset, which includes phylogenetic classification into 20 families, were randomly sampled to a maximum of 10 TerL sequences per family, for a total of 200. TerL hits in the IMG/VR dataset were single-linkage clustered by pairwise sequence identity with a threshold (17.5%) chosen to result in exactly 30 clusters, then one random sequence was taken from each cluster. The final TerL set consists of 238 sequences: 8 identifiable TerL homologs from our 20 Millard genomes containing candidate introns, 200 representative TerL homologs from phylogenetically annotated intron-negative Millard genomes, and 30 representative TerL homologs from the broader but phylogenetically unannotated IMG/VR genomes. These sets of RVT and TerL protein sequences were then aligned with MAFFT v7.525 (27), and phylogenetic tree inference was done with IQ-TREE2 v2.3.0 with the “model finder plus” flag (28; 29) and 1000 bootstrap replicates. Unrooted trees were visualized with iToL (30). For all pairwise percent identity (PID) values, the denominator is the shorter of the two unaligned sequence lengths.

Secondary structure predictions were done using RNAfold (31) and mfold (32) and visualized using rnacanvas (33). Genomic context of introns was plotted using LoVis4u (34).

## Results and Discussion

The Rfam database of conserved RNA structure families includes curated multiple sequence alignments and profile SCFGs for 4000+ conserved structural RNAs (18). Rfam includes eight models of fragments of the group II intron consensus, built from alignments of known eukaryotic organellar and bacterial group II intron sequences. The most conserved region of the intron is the 3^′^ D5/D6 region of the intron, Rfam model RF00029. The 5^′^ end of the intron, domains D1-D4, varies widely across group II intron classes, and is represented in Rfam by a set of seven models grouped into an Rfam “clan”, CL00102. For each complete group II intron we expect to find one or two hits: a D1-D4 hit to one of the seven 5^′^ end models, and a D5/D6 hit a few hundred nucleotides or around 1 kb downstream, for an orfless and ORF-containing intron, respectively.

Available phage genome sequences have grown rapidly in recent years, collected in different places. We started with a well-curated phage genome dataset provided by the Millard lab, comprising 29,015 phage genomes totaling 1.77 Gb. Using the Infernal search program cmsearch, we found 27 hits to the D1-D4 models and 23 to the D5/D6 model in the Millard phage genomes with E-value *<* 0.01.

Looking at these hits, we removed four genomes (MK448731, MK448731, MK448781, MK448888) where putative intron regions were identical to other representative genomes; one genome (NC 030940) where a group II intron occurs just before the prophage integration site and appears to be misannotated as being within the prophage; and three scaffolds which appear to be integrated conjugative elements (MT836071, MT836602, MT836027). We kept both of two other group II introns that are identical in sequence (in NC 031039 and NC 043027), because both phage (AR9 and PBS1) have been well studied, and the host RNA polymerase gene and a downstream group I intron in it are not identical.

After these removals, we had a set of 20 putative group II introns in phage genomes in the Millard database (Figure 2). Of these, 12 have both D1-D4 and D5/D6 hits, 4 have D1-D4 and no D5/D6, and 4 with only a D5/D6 hit. One intron hit, OQ555808, contained two D1D4 hits: to model group-II-D1D4-3, followed by group-II-D1D4-1. Upstream and downstream hits, when both present, were always within 2 kb of each other, as expected for typical group II intron length.

**Fig. 2:**
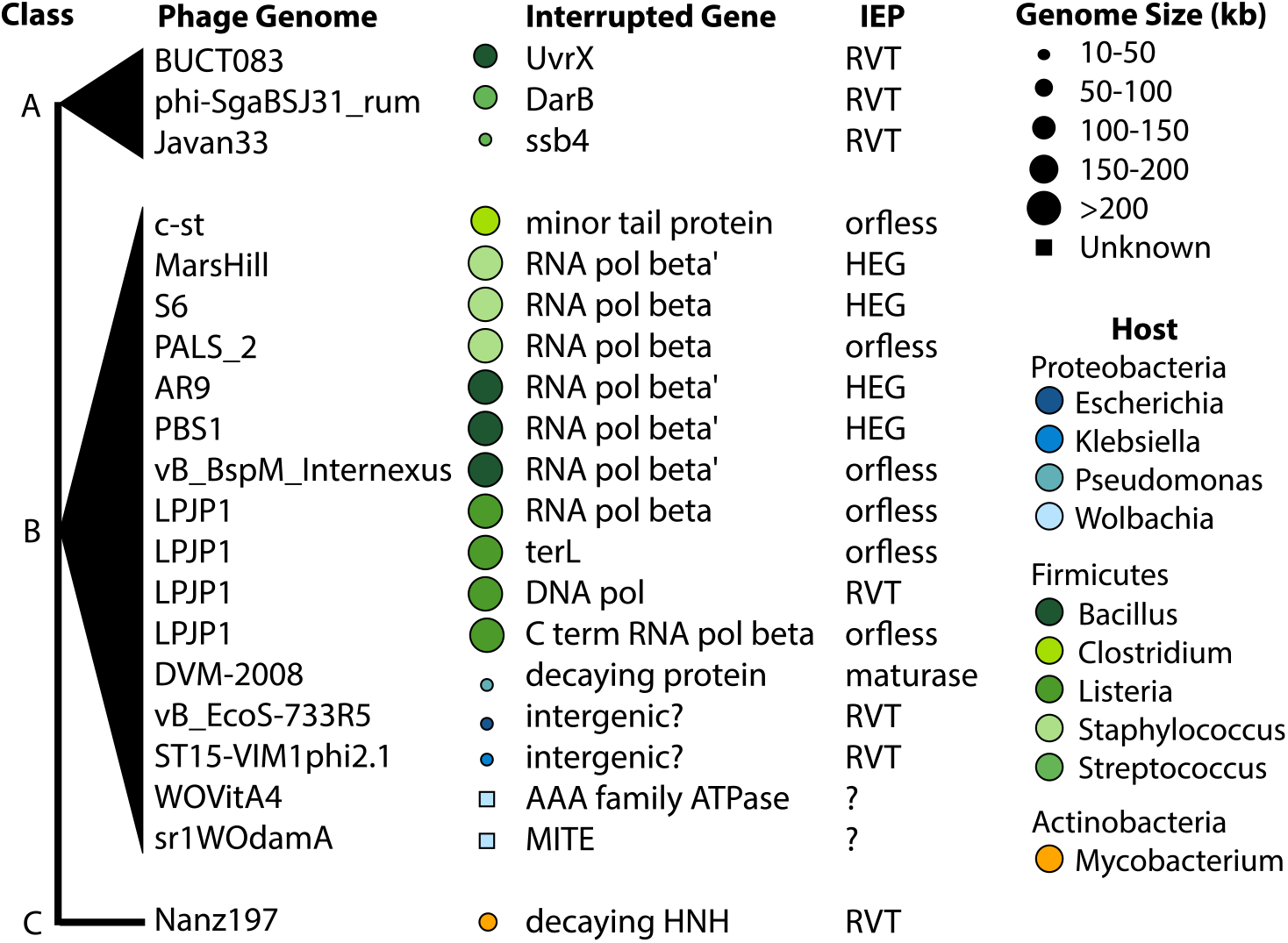
Phage group II introns. Genome size is indicated by size of circle to right of genome name, and host bacterial taxon by color.Two genomes were incomplete contigs (WOVitA4 and sr1WOdamA) and thus their genome size bubble is denoted as a box (for unknown). Introns are sorted by class: IIA, IIB, or IIC-type introns.

We then used cmsearch with the same eight Rfam models to search the much larger IMG/VR metagenomic viral database of 15.7 million contigs and found putative intron hits in 9229 genomes: 6205 hits to the D1-D4 models and 6703 to the D5/D6 model with E-value *<* 0.01 (Supplementary Table S2). The proportion of intron-containing genomes was approximately the same between Millard (20 of 29015 = 0.07%) and IMG/VR (9229 of 15722824 = 0.06%). The IMG/VR phage hits included an unusual one worth noting: contig IMGVR UViG 3300011577 000009 is annotated as a high-confidence *Inovirus*, a single-stranded DNA phage. The group II intron mobility mechanism targets dsDNA, and no ssDNA genomes with group II introns have been described previously. We used hmmscan and Pfam to confirm that this phage genome contig contains homologs distinctive of *Inoviridae*, a Zot domain and a replication G2P protein. The intron itself appears to be complete, intact, and contains a clear RVT IEP. We imagine that an ssDNA virus could be infected by a group II intron during replication, when it exists in a dsDNA intermediate.

We searched with additional profile SCFGs we built from the newly identified phage introns, to account for the possibility that the existing Rfam models might imperfectly represent their sequence diversity. We made a new profile SCFG of the alignment of 6703 D5/D6 hits in IMG/VR hits (using the cmsearch -A), then re-searched IMG/VR using this alignment. This identified hits in another 680 IMG/VR contigs besides the previous 9229. We made two full-length group II intron models, one of the 3 type IIA and the other of the 16 type IIB introns (Figure 2), starting from our inferred structures for ON107264 and NC 007581, respectively. These models identified 711 and 97 new hits in IMG/VR. These results indicated that more refined and/or phage-specific models could reveal some additional group II introns, but the gains from the effort seemed incremental. None of the three phage-specific models identified additional candidate introns in the genomes in the Millard dataset.

We used three additional lines of evidence, in addition to significant Infernal search results, to improve our confidence in the set of 20 putative group II introns. First, we sought to confirm that candidates are indeed in intervening sequences. We stitched together flanking exons to infer the host gene protein sequence, then used this to identify intronless homologs in other phage. We also used the alignment to the closest intronless homolog to help define the 5^′^ and 3^′^ intron boundaries (Supplementary Table S1). The 3’ splice site is generally closely determined by the hit to the conserved D5/D6 model, and the 5’ splice site occurs at a conserved GWYRG site (35; 4) in the upstream vicinity of the hits to Rfam D1-D4 models. Fourteen of the 20 candidate introns are in intervening sequences in host genes where we could identify and align to an intronless homolog. Of the remaining six, two appear to be intergenic (ON470608, MK448228), one appears in a heavily decayed pseudogene (EU982300), two are incomplete intron sequences with start or ends outside of the contig boundaries (HQ906664, KY695241), and one is downstream of a rho-independent terminator in a typical position for bacterial group IIC introns (OQ555808; Supplementary Figure 1) (36).

Second, we identified a set of consensus pseudoknotted base-pairing interactions critical for group II intron function using secondary structure prediction tools and manual curation. Because profile SCFG algorithms do not model pseudoknots, these additional base-pairing interactions provide independent evidence in support of a group II intron call. Specifically, we looked for pseudoknot base-pairing interactions EBS1:IBS1, EBS2:IBS2, *α* : *α*^′^, *β* : *β*^′^, *δ* : *δ*^′^, *ϵ* : *ϵ*^′^, *γ* : *γ*^′^, and also for tertiary non-canonical pairing interactions *λ* : *λ*^′^, *κ* : *κ*^′^, *ζ* : *ζ*^′^. Figure 3 shows an example structure of one of the phage introns, with these interactions annotated. We used the region around *λ* − *ϵ*^′^ to classify the group II introns by subtype, where A has an 11 nt loop with consensus AGC, B is a 4 nt bulge with consensus AARC, and C contains a 7-12 nt loop with consensus AGG (2; 35). To confirm the class affiliation of each intron, we use primary sequence similarity to previously classified bacterial introns using the Zimmerly and Candales database (37). Examples of all three classes were identified (Figure 2).

**Fig. 3:**
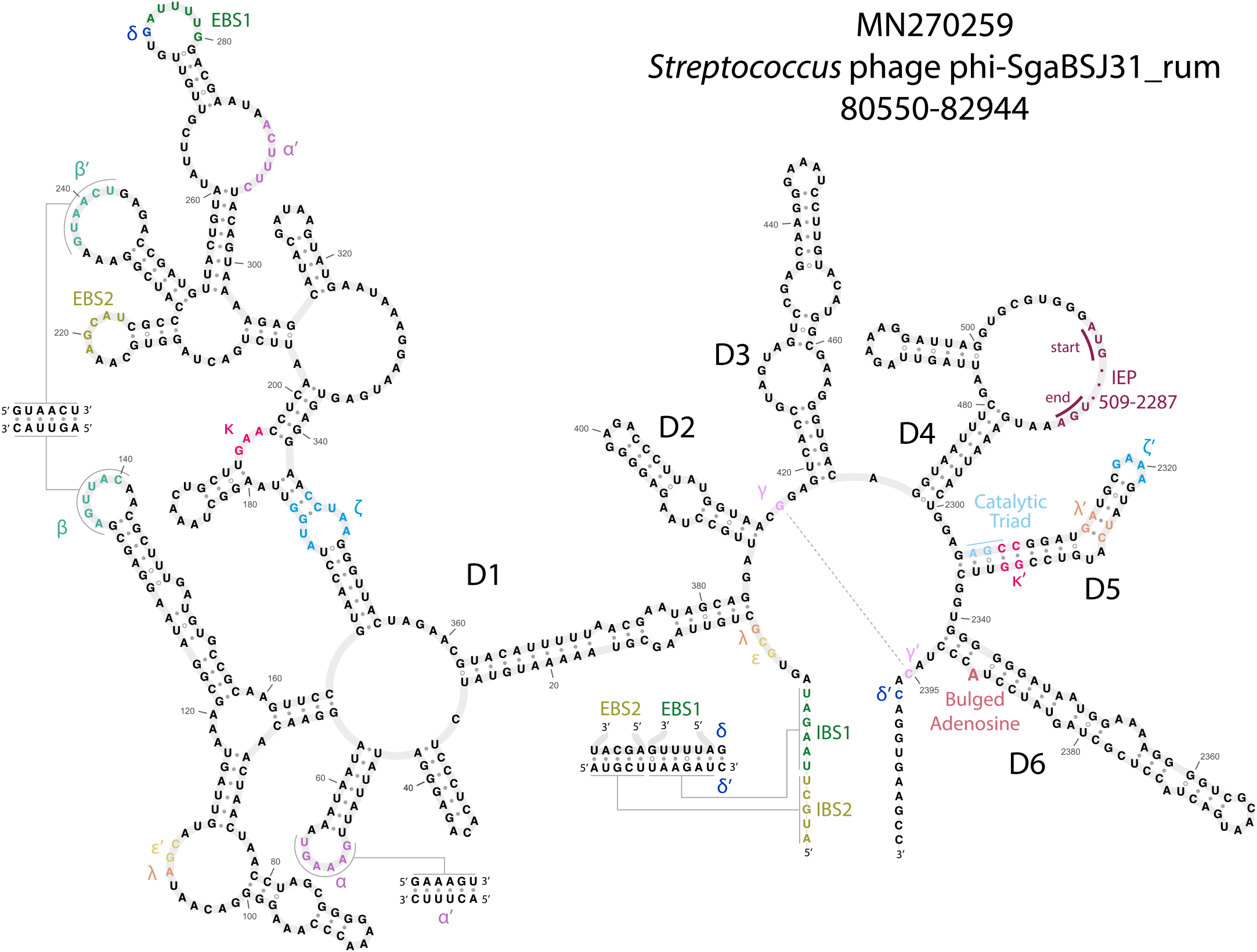
Secondary structure example. Manually curated predicted secondary structure of the group IIA intron in a DarB-like antirestriction/SNF2 helicase gene in *Streptococcus* phage phi-SgaBSJ31_rum.

Lastly, we analyzed ORFs encoded by the 20 group II introns. We translated the regions within the intron to identify a total of 7 RVT IEP and 4 homing endonucleases (HEGs) (Figure 2; Supplementary Table S1). Many types of reverse transcriptases occur in bacterial and phage genomes, including retrons, CRISPR-Cas systems, diversity generating retroelements, and phage defense systems (26; 38; 39; 40). The seven RVT IEPs were generally misannotated in GenBank files, often as “retron-type” reverse transcriptases. Phylogenetic tree inference placed all seven IEP RVTs within the clade of known group II intron IEPs to the exclusion of other known RVT clades including retrons (Figure 4A). All seven RVT IEPs also have the “X” maturase domain characteristic of a group II IEP (Figure 4B).

**Fig. 4:**
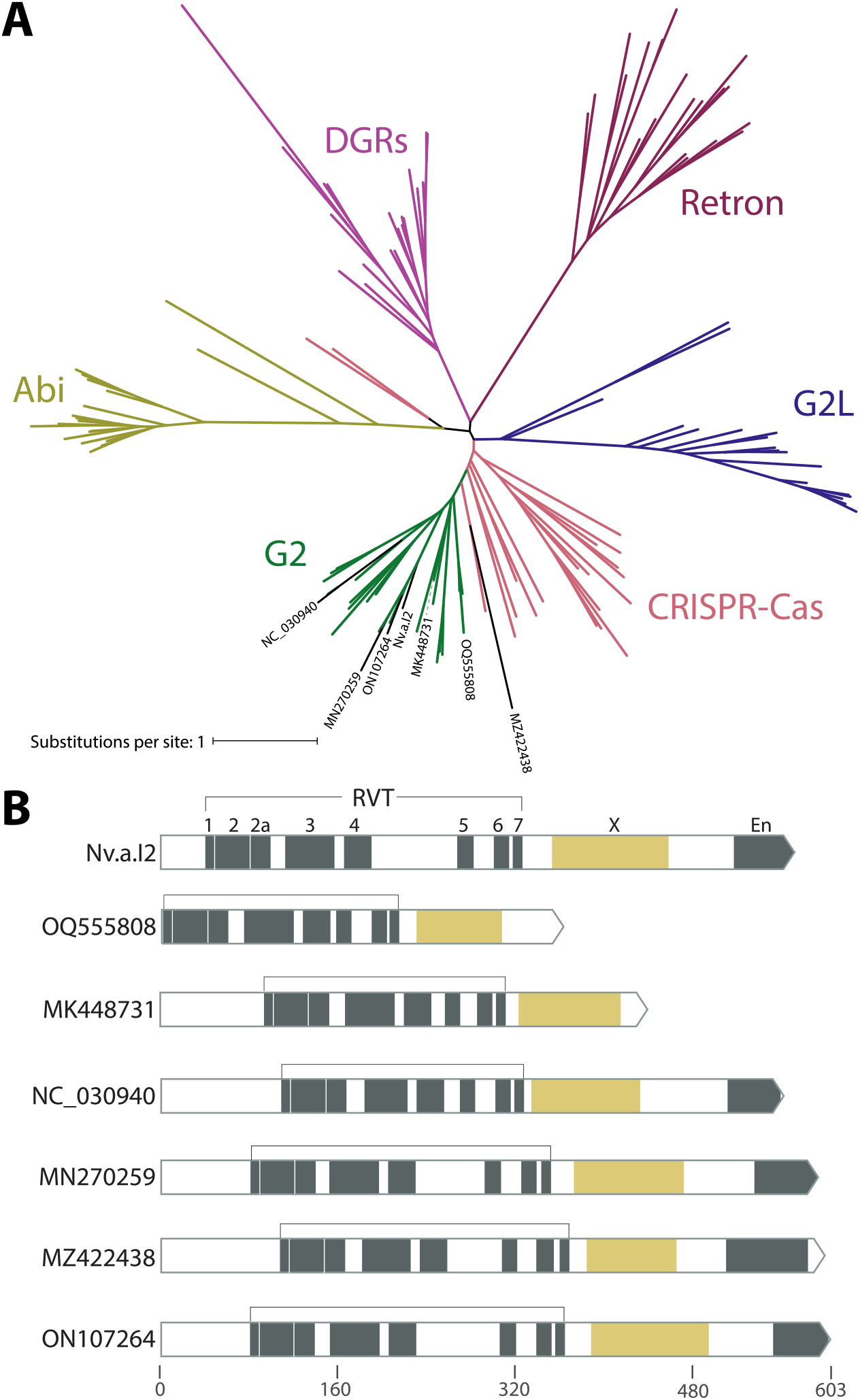
Reverse transcriptase phylogeny. **A**. Unrooted tree of reverse transcriptases found in various genetic elements in bacteria and phage. RVT domains of known group II intron IEPs are shown in green, and RVT domains of phage group II IEPs identified in this paper labeled in black. Abi = abortive infection defense systems; DGRs = diversity generating elements; G2L = “group II-like”. **B**. Domain structure of phage group II IEPs includes the maturase “X” domain characteristic of group II IEPs, and the frequent presence of an endonuclease domain. For comparison, domain structure of a known group II IEP is shown at top (Nv.a.I2, a bacterial group IIA intron in *Novosphingobium aromaticivorans* (37)).

Six introns are orfless, but this does not necessarily imply that they are immobile. An orfless phage group II intron may retain mobility if another group II intron in the same cell provides an IEP in *trans* (41). For example, one phage genome (LPJP1, a *Listeria* jumbophage) contains four group II introns, one of which encodes an RVT IEP and three of which are orfless (Figure 2). This seems likely to be a case of IEP borrowing.

We looked closely at the four intron ORFs that appear to encode homing endonucleases instead of IEPs. HEGs are more typical of mobile group I introns, and indeed we find additional homologs of these four HEGs elsewhere in the same phage genomes and mostly in group I introns, including a case of HEG-containing group I and group II introns in the same host gene (Figure 5A)). HEG-based mobility works by the HEG making a specific dsDNA cleavage at the intron insertion site in an intronless allele, which is then a substrate for double-strand break repair recombination using the intact intron-plus allele as the repair template. Unlike the group II intron retrohoming mechanism, HEG-mediated mobility is a DNA-level event independent of RNA catalysis, so HEGs are found in many types of mobile sequences, including group I introns, bulge-helix-bulge introns, and inteins, and free-standing mobile intergenic HEGs are also common (42). A few cases of group II introns with HEGs have been observed before, for two of the most abundant HEG families, the LAGLIDADG and GIY-YIG HEGs (43; 44). These four phage group II HEGs are not closely related to known HEG families.

**Fig. 5:**
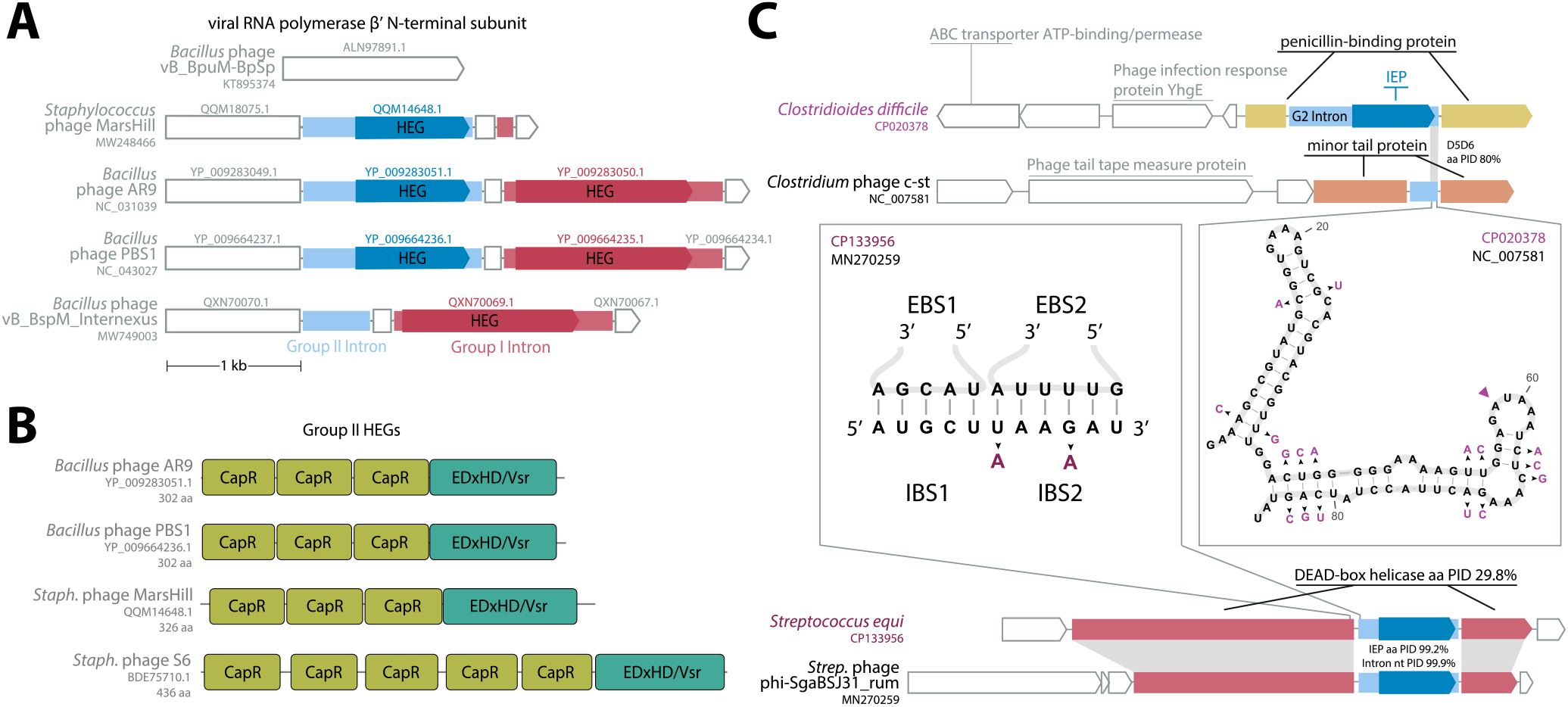
Homing endonuclease genes found in four introns; close bacterial relatives found for two. **A**. Phage genes for the virion RNA polymerase *β*^′^ N-terminal subunit are invaded by both group I and group II introns with homologous HEGs. Group I introns are shown in red, with their encoded HEGs shown in darker red. Group II introns are shown in light blue, with their encoded HEG in darker blue. **B**. Four group II introns (including three shown in panel A) encode a large new outgroup of HEGs which consist of several domains distant related to Pfam CapR domains (light green) followed by a putative nuclease domain distantly homologous to EDxHD/Vsr homing enconucleases (teal). **C**. Two phage group II introns have closely related bacterial homologs. Phage loci are labeled in black; host loci are labeled in purple. One case (top diagram and right panel) is a possible retroposition event between unrelated host genes. The other case (bottom diagram and left panel) involves homologous host genes in the phage and bacterial genomes. The structure inset (right panel) shows the D5/D6 region of the phage intron, and substitutions in the bacterial homolog are shown in purple.

HMMER profile analysis and AlphaFold3 structure prediction (45) identifies a conserved domain structure with three to five ∼60aa domains that are distantly related to Pfam models CapR, DUF4379, and DUF723, followed by a C-terminal ∼100aa domain distantly related to the known EDxHD/Vsr HEG endonuclease domain (46) (Figure 5B). The CapR-like domain is likely to be a DNA-binding domain that confers additional sequence specificity onto the HEG endonuclease domain, as seen with several known families of auxiliary “NUMODs” (nuclease-associated modular DNA-binding domains) (47). Profile HMMs for these CapR-like and EDxHD/Vsr-like domains find thousands of hits in UniProt, many of which are unannotated hypothetical proteins, so this appears to be a large unrecognized outgroup of the EDxHD/Vsr HEGs.

One way for a phage genome to acquire group II introns is by retroposition from the host bacterial genome. This would be more likely to occur for lysogenic phage with an integrated prophage stage in their life cycle, as opposed to lytic phage. We searched for bacterial group II introns similar to our phage introns and identified two examples (Figure 5C). In both cases, the phage intron with an identifiable bacterial sequence homolog in in a phage annotated as lysogenic prophage. For the group II intron in prophage c-st, we found a diverged host intron (about 80% identical in D5/D6) in a nonhomologous host gene. For the intron in prophage phi-SgaBSJ31 rum, we found a near-identical intron in a helicase gene remotely homologous to the phage host gene, a DEAD-box helicase annotated as a DarB antirestriction system. The immediate exonic flanking regions of the phage and bacterial introns are similar, with 2 substitutions in IBS2, one of which conserves pairing, and the of other which is the dispensable first position of IBS2 (since EBS2:IBS2 can be as short as 4 bp).

We sought to determine whether group II introns are widespread across different types of phage, or if they are confined to a particular phylogenetic clade. While there is no universal phylogenetic marker for phage, all twenty of our examples from the Millard dataset (and most of the IMG/VR hits) are within *Caudoviricetes*, for which the terminase large subunit TerL is often used for taxonomic classification (48; 49). A phylogenetic tree inferred from an alignment of terminase TerL protein sequences from both intron-plus and intron-minus representatives across all families represented in the Millard database shows that group II introns are dispersed across *Caudoviricetes* phage phylogeny (Figure 6).

**Fig. 6:**
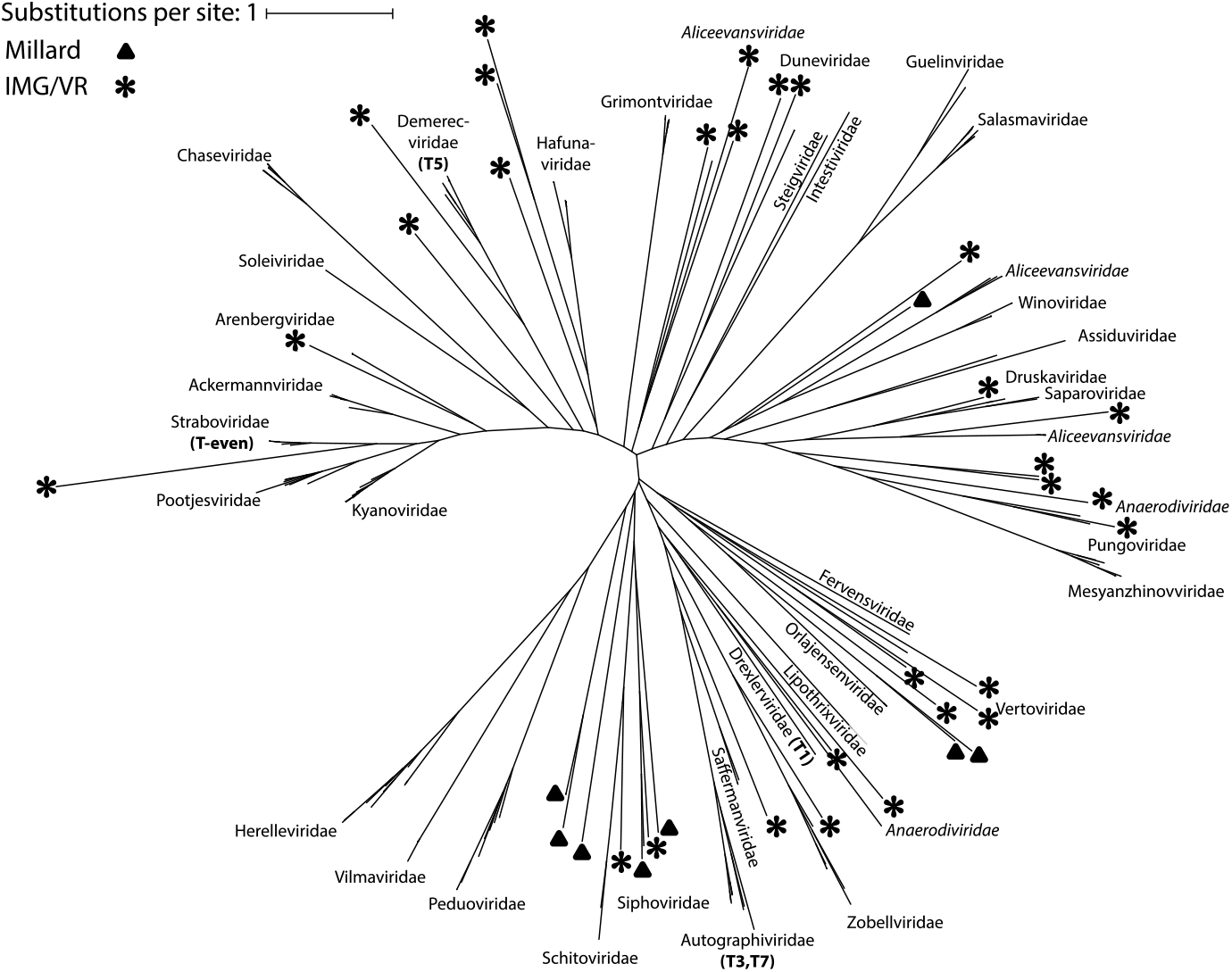
Phylogenetic distribution of phage group II introns. Unrooted tree using an alignment of terminase large subunit (TerL) protein sequences, with phage families labeled. Italics indicating a polyphyletic family in the tree. Taxa containing group II introns are denoted with a black triangle (Millard dataset) or star (IMG/VR).

Our results extend and generalize two previously observed cases of group II introns in phage genomes (8; 9), neither of which had noted that group II introns had been believed to be absent from phage (7). Group II introns are relatively prevalent and occur in a wide variety of phage genomes. These introns are usually not annotated in GenBank, and group II RVT IEPs are typically being misannotated as other types of reverse transcriptase containing elements, suggesting that standard phage genome annotation pipelines could be improved to look for these elements.

Besides phage, another place that group II introns are thought to be absent is eukaryotic nuclear genomes. Using the latest accelerated versions of Infernal, systematic searches of all eukaryotic nuclear genomes with group II intron profile SCFGs are feasible.

## Competing interests

No competing interest is declared.

## Acknowledgments

Thank you to Prof. Aaron Robart for intron classification discussions. We thank the FAS Research Computing Group at Harvard University for our high-performance computational resources on the Cannon cluster. We gratefully acknowledge the US Department of Energy Joint Genome Institute (https://www.jgi.doe.gov/) and its the user community for our use of the IMG/VR dataset.

## Funding

This work was supported the National Science Foundation [Graduate Research Fellowship to L.N.M], the National Institute of Health [Harvard Molecular Biophysics training grant T32GM008313 and R01-HG009116 to S.R.E.], and the Howard Hughes Medical Institute.

## Notes

https://doi.org/10.5281/zenodo.15490104

### Competing Interest Statement

The authors have declared no competing interest.

## References

1. Lambowitz AM, Belfort M. Mobile Bacterial Group II Introns at the Crux of Eukaryotic Evolution. Microbiology spectrum. 2015;3:MDNA3–00502014.

2. Michel F, Jacquier A, Dujon B. Comparison of fungal mitochondrial introns reveals extensive homologies in RNA secondary structure. Biochimie. 1982;64:867–81.

3. Michel F, Dujon B. Conservation of RNA secondary structures in two intron families including mitochondrial-, chloroplast- and nuclear-encoded members. The EMBO Journal. 1983;2:33–8.

4. Zimmerly S, Semper C. Evolution of group II introns. Mobile DNA. 2015;6:7.

5. Miura MC, Nagata S, Tamaki S, Tomita M, Kanai A. Distinct Expansion of Group II Introns During Evolution of Prokaryotes and Possible Factors Involved in Its Regulation. Frontiers in Microbiology. 2022;13.

6. Toro N, Jiménez-Zurdo JI, García-Rodríguez FM. Bacterial group II introns: not just splicing. FEMS Microbiology Reviews. 2007;31:342–58.

7. Edgell DR, Belfort M, Shub DA. Barriers to Intron Promiscuity in Bacteria. Journal of Bacteriology. 2000;182:5281–9.

8. Lavysh D, Sokolova M, Minakhin L, Yakunina M, Artamonova T, Kozyavkin S, et al. The genome of AR9, a giant transducing Bacillus phage encoding two multisubunit RNA polymerases. Virology. 2016;495:185–96.

9. Korn AM, Hillhouse AE, Sun L, Gill JJ. Comparative Genomics of Three Novel Jumbo Bacteriophages Infecting Staphylococcus aureus. Journal of Virology. 2021;95:10.1128/jvi.02391-20.

10. Hausner G, Hafez M, Edgell DR. Bacterial group I introns: mobile RNA catalysts. Mobile DNA. 2014;5:8.

11. Dai L, Zimmerly S. Compilation and analysis of group II intron insertions in bacterial genomes: evidence for retroelement behavior. Nucleic Acids Res. 2002;30:1091–102.

12. Lang BF, Laforest MJ, Burger G. Mitochondrial introns: a critical view. Trends in Genetics. 2007;23:119–25.

13. García-Rodríguez FM, Neira JL, Marcia M, Molina-Sánchez MD, Toro N. A group II intron-encoded protein interacts with the cellular replicative machinery through the -sliding clamp. Nucleic Acids Res. 2019;47:7605–17.

14. Toor N, Hausner G, Zimmerly S. Coevolution of group II intron RNA structures with their intron-encoded reverse transcriptases. RNA. 2001;7:1142–52.

15. Eddy SR, Durbin R. RNA sequence analysis using covariance models. Nucleic Acids Res. 1994;22:2079–88.

16. Sakakibara Y, Brown M, Hughey R, Mian IS, Sjölander K, Underwood RC, et al. Stochastic context-free grammars for tRNA modeling. Nucleic Acids Res. 1994;22:5112–20.

17. Nawrocki EP, Eddy SR. Infernal 1.1: 100-fold faster RNA homology searches. Bioinformatics. 2013;29:2933–5.

18. Kalvari I, Nawrocki EP, Ontiveros-Palacios N, Argasinska J, Lamkiewicz K, Marz M, et al. Rfam 14: expanded coverage of metagenomic, viral and microRNA families. Nucleic Acids Res. 2021;49:D192–200.

19. Nawrocki EP, Jones TA, Eddy SR. Group I introns are widespread in archaea. Nucleic Acids Res. 2018;46:7970–6.

20. Cook R, Brown N, Redgwell T, Rihtman B, Barnes M, Clokie M, et al. INfrastructure for a PHAge REference Database: Identification of Large-Scale Biases in the Current Collection of Cultured Phage Genomes. PHAGE. 2021;2:214–23.

21. Camargo AP, Nayfach S, Chen IMA, Palaniappan K, Ratner A, Chu K, et al. IMG/VR v4: an expanded database of uncultivated virus genomes within a framework of extensive functional, taxonomic, and ecological metadata. Nucleic Acids Res. 2023;51:D733–43.

22. Schwengers O, Jelonek L, Dieckmann MA, Beyvers S, Blom J, Goesmann A. Bakta: rapid and standardized annotation of bacterial genomes via alignment-free sequence identification. Microbial Genomics. 2021;7:000685.

23. Bouras G, Nepal R, Houtak G, Psaltis AJ, Wormald PJ, Vreugde S. Pharokka: a fast scalable bacteriophage annotation tool. Bioinformatics. 2023;39:btac776.

24. Eddy SR. Accelerated Profile HMM Searches. PLOS Computational Biology. 2011;7:e1002195.

25. Mistry J, Chuguransky S, Williams L, Qureshi M, Salazar GA, Sonnhammer ELL, et al. Pfam: The protein families database in 2021. Nucleic Acids Res. 2021;49:D412–9.

26. Toro N, Martínez-Abarca F, Mestre MR, González-Delgado A. Multiple origins of reverse transcriptases linked to CRISPR-Cas systems. RNA Biology. 2019;16:1486–93.

27. Katoh K, Standley DM. MAFFT Multiple Sequence Alignment Software Version 7: Improvements in Performance and Usability. Molecular Biology and Evolution. 2013;30:772–80.

28. Minh BQ, Schmidt HA, Chernomor O, Schrempf D, Woodhams MD, von Haeseler A, et al. IQ-TREE 2: New Models and Efficient Methods for Phylogenetic Inference in the Genomic Era. Molecular Biology and Evolution. 2020;37:1530–4.

29. Kalyaanamoorthy S, Minh BQ, Wong TKF, von Haeseler A, Jermiin LS. ModelFinder: fast model selection for accurate phylogenetic estimates. Nature Methods. 2017;14:587–9.

30. Letunic I, Bork P. Interactive Tree of Life (iTOL) v6: recent updates to the phylogenetic tree display and annotation tool. Nucleic Acids Res. 2024;52:W78–82.

31. Gruber AR, Lorenz R, Bernhart SH, Neuböck R, Hofacker IL. The Vienna RNA Websuite. Nucleic Acids Res. 2008;36:W70–4.

32. Zuker M. Mfold web server for nucleic acid folding and hybridization prediction. Nucleic Acids Res. 2003;31:3406–15.

33. Johnson PZ, Simon AE. RNAcanvas: interactive drawing and exploration of nucleic acid structures. Nucleic Acids Res. 2023;51:W501–8.

34. Egorov AA, Atkinson GC. LoVis4u: a locus visualization tool for comparative genomics and coverage profiles. NAR Genom Bioinform. 2025;7:lqaf009.

35. Michel F, Kazuhiko U, Haruo O. Comparative and functional anatomy of group II catalytic introns — a review. Gene. 1989;82:5–30.

36. Robart AR, Seo W, Zimmerly S. Insertion of group II intron retroelements after intrinsic transcriptional terminators. PNAS. 2007;104:6620–5.

37. Candales MA, Duong A, Hood KS, Li T, Neufeld RAE, Sun R, et al. Database for bacterial group II introns. Nucleic Acids Res. 2012;40:D187–90.

38. Wilkinson ME, Li D, Gao A, Macrae RK, Zhang F. Phage-triggered reverse transcription assembles a toxic repetitive gene from a noncoding RNA. Science. 2024;0:eadq3977.

39. Tang S, Conte V, Zhang DJ, Žedaveinytė R, Lampe GD, Wiegand T, et al. De novo gene synthesis by an antiviral reverse transcriptase. Science. 2024;0:eadq0876.

40. González-Delgado A, Mestre MR, Martínez-Abarca F, Toro N. Prokaryotic reverse transcriptases: from retroelements to specialized defense systems. FEMS Microbiology Reviews. 2021;45:fuab025.

41. Meng Q, Wang Y, Liu XQ. An intron-encoded protein assists RNA splicing of multiple similar introns of different bacterial genes. The Journal of Biological Chemistry. 2005;280:35085–8.

42. Stoddard BL. Homing endonucleases from mobile group I introns: discovery to genome engineering. Mobile DNA. 2014;5:7.

43. Mullineux ST, Costa M, Bassi GS, Michel F, Hausner G. A group II intron encodes a functional LAGLIDADG homing endonuclease and self-splices under moderate temperature and ionic conditions. RNA. 2010;16:1818–31.

44. Lambowitz AM, Zimmerly S. Group II Introns: Mobile Ribozymes that Invade DNA. Cold Spring Harbor Perspectives in Biology. 2011;3:a003616.

45. Abramson J, Adler J, Dunger J, Evans R, Green T, Pritzel A, et al. Accurate structure prediction of biomolecular interactions with AlphaFold 3. Nature. 2024;630:493–500.

46. Dassa B, London N, Stoddard BL, Schueler-Furman O, Pietrokovski S. Fractured genes: a novel genomic arrangement involving new split inteins and a new homing endonuclease family. Nucleic Acids Res. 2009;37:2560–73.

47. Sitbon E, Pietrokovski S. New types of conserved sequence domains in DNA-binding regions of homing endonucleases. Trends in Biochemical Sciences. 2003;28:473–7.

48. Yutin N, Benler S, Shmakov SA, Wolf YI, Tolstoy I, Rayko M, et al. Analysis of metagenome-assembled viral genomes from the human gut reveals diverse putative CrAss-like phages with unique genomic features. Nature Communications. 2021;12:1044.

49. Yutin N, Tolstoy I, Mutz P, Wolf YI, Krupovic M, Koonin EV. Jumping DNA polymerases in bacteriophages. bioRxiv. 2024:2024.04.26.591309.

50. D’Souza LM, Zhong J. Mutations in the Lactococcus lactis Ll.LtrB group II intron that retain mobility in vivo. BMC Molecular Biology. 2002;3:17.

51. Belfort M, Lambowitz AM. Group II Intron RNPs and Reverse Transcriptases: From Retroelements to Research Tools. Cold Spring Harbor Perspectives in Biology. 2019;11:a032375.

