## Supplementary Figure 1 for "Prevalence of Group II Introns in Phage Genomes"

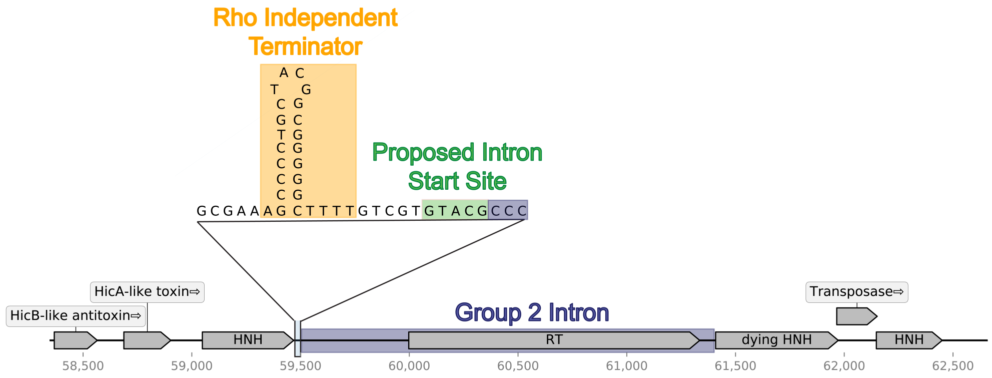


**Figure 1: A phage group IIC intron.** Group IIC introns typically insert 4-8 nucleotides downstream of a rho-independent transcriptional terminator, rather than in the coding region of their host gene. One of our phage group II introns is a type IIC intron, and we can identify a rho-independent terminator 5 nucleotides upstream of its insertion site.
